# A large-scale multi-omics study of cortical miRNA dysregulation in Alzheimer’s disease

**DOI:** 10.64898/2026.11.16.767087

**Authors:** Ehsan Pishva, Morteza Kouhsar, M. Muaaz Aslam, Marcel Schilling, Joshua Harvey, Darren Soanes, Jennifer Imm, Grace Thomson, Giulia Pegoraro, Luke Weymouth, Lachlan F. MacBean, Gregory Wheildon, Valerija Dobricic, Assia Tiane, Melissa Schepers, Jarno Koetsier, Valentin Laroche, Tim Vanmierlo, Daniel van den Hove, Claire Troakes, Safa Al-Sarraj, Laura Parkkinen, Jonathan Mill, Byron Creese, Adam R. Smith, Lars Bertram, Katie Lunnon

**Author notes:** Corresponding authors: Ehsan Pishva: Department of Psychiatry and Neuropsychology, Mental Health and Neuroscience Research Institute (MHeNs), Maastricht University, Maastricht, The Netherlands, Katie Lunnon: Department of Clinical and Biomedical Sciences, Faculty of Health and Life Sciences, Exeter University, Exeter, UK.

## Abstract

MicroRNAs regulate gene expression, but their roles in Alzheimer’s disease pathology and downstream molecular changes remain incompletely characterized. We integrated cortical small RNA-sequencing data from seven datasets across six postmortem cohorts comprising 1,738 individuals with transcriptomic, proteomic, genetic, and single-nucleus data. Meta-analysis of 405 microRNAs identified 16 associated with advanced and four with intermediate neurofibrillary tangle pathology, with miR-132-3p significant in both comparisons. Five microRNAs (miR-132-3p, miR-212-3p, miR-212-5p, miR-146b-5p, miR-3200-3p) showed evidence of mediating associations between target mRNA and protein expression and tau pathology. Consensus co-expression network analysis linked the miR-132/212 family to opposing neuronal/synaptic and oligodendrocyte/myelination-related programmes. Mediated targets were enriched among tau-associated differentially expressed genes across glial cell types, particularly astrocytes. Inferred cell-type regulatory networks further highlighted miR-132-3p and miR-146b as recurrent hubs. These findings prioritize a restricted set of cortical microRNAs and associated neuronal and glial molecular programmes for mechanistic investigation in Alzheimer’s disease.

## Main

MicroRNAs (miRNAs) are a highly conserved class of small non-coding RNAs that regulate post-transcriptional gene expression and are key regulators of complex traits and diseases. Growing evidence suggests that miRNA dysregulation contributes to the molecular mechanisms underlying Alzheimer’s disease (AD), including amyloid processing, tau pathology, neuroinflammation, and synaptic dysfunction^1–7^. A single miRNA can regulate numerous mRNA targets, while individual transcripts may be targeted by multiple miRNAs, making it inherently challenging to disentangle the contribution of specific miRNAs to AD-associated molecular networks. Although numerous studies have reported altered miRNA expression in AD^8–10^, genome-wide studies of human post-mortem brain tissue have generally been underpowered and have yielded inconsistent findings. Furthermore, comprehensive multi-omic studies integrating miRNA, transcriptomic, and proteomic data at scale remain scarce. Consequently, it remains unclear which miRNA alterations are robustly associated with AD pathology and how these changes influence downstream molecular pathways involved in neurodegeneration. This has hindered the identification of key miRNA-mediated regulatory mechanisms and their translation into potential therapeutic strategies.

Here, we conducted a large-scale multi-omic investigation of cortical miRNA dysregulation in AD, integrating matched genome-wide miRNA, mRNA, protein, and genetic data across multiple independent postmortem brain cohorts. Through this approach, we identified miRNAs robustly associated with AD pathology, investigating their relationships with downstream gene and protein expression, and characterized the biological pathways and cellular contexts in which they may act. Together, these analyses provide a comprehensive systems-level characterization of miRNA-mediated regulatory networks in AD cortical brain tissue.

## Results

### Meta-analysis of miRNA expression associated with tau pathology

To identify cortical miRNAs associated with progressive neurofibrillary tangle (NFT) pathology, we performed a cross-cohort meta-analysis of small RNA-sequencing (small RNA-seq) data from seven datasets generated across six independent postmortem brain cohorts comprising 1,738 individuals (**Fig. 1**, **Supplementary Table 1**). Analyses were restricted to the 405 miRNAs consistently detected across all cohorts and were performed separately for advanced pathology (n = 1,178, seven datasets; Braak V-VI: n = 690, Braak 0-II: n = 488) and intermediate pathology (n = 1,048, five datasets; Braak III-IV: n = 560, Braak 0-II: n = 488) because of the unequal distribution of intermediate-stage samples across all datasets.

**Fig. 1:**
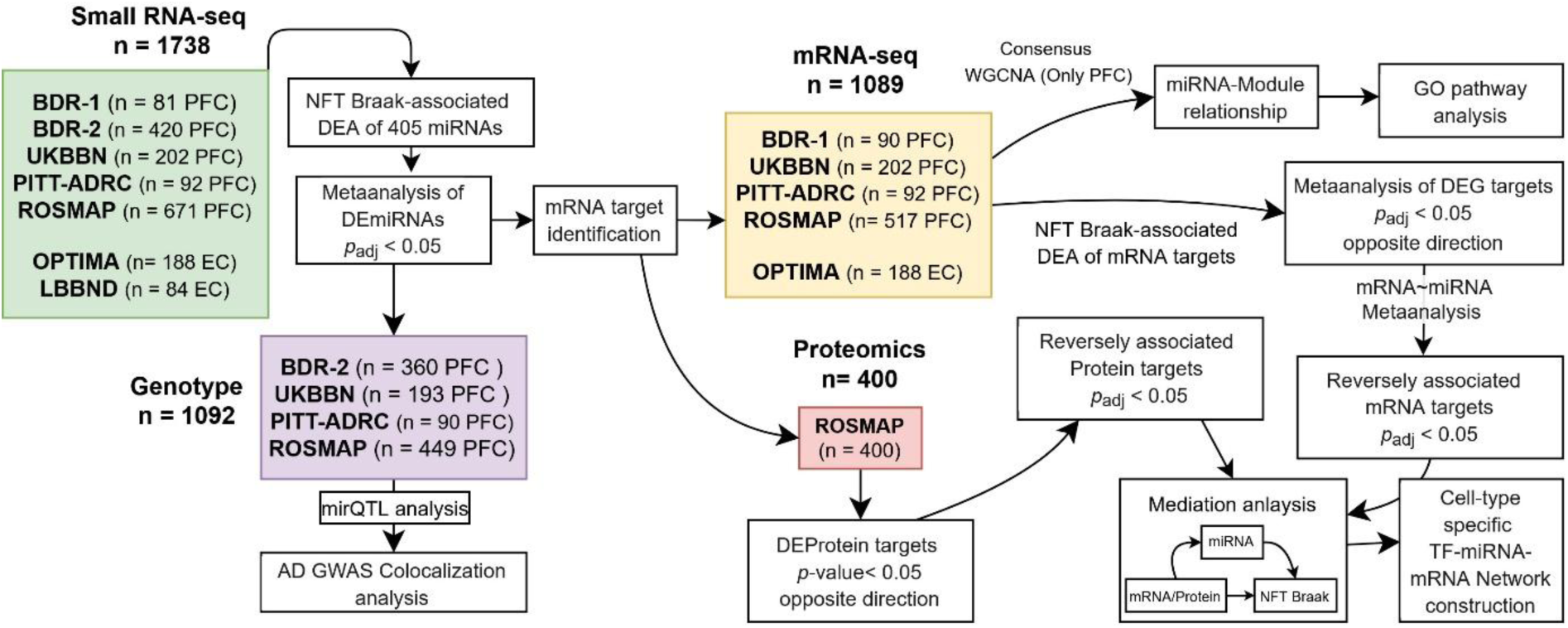
Overview of study design. Seven small RNA-sequencing datasets from six independent human brain cohorts were analyzed to identify miRNAs associated with Braak stage through differential expression analysis and meta-analysis. Downstream analyses included target identification, miRNA quantitative trait locus (miQTL) mapping, and Bayesian colocalization with Alzheimer’s disease (AD) genome-wide association signals. Integration with transcriptomic and proteomic data identified inversely associated mRNA and protein targets. Consensus weighted gene co-expression network analysis (WGCNA) linked miRNAs to gene modules and enriched Gene Ontology (GO) pathways. Mediation analyses assessed relationships among miRNAs, target expression, and neurofibrillary (NFT) tangle pathology, complemented by cell-type-specific transcription factor (TF)-miRNA–mRNA network construction. Cohort lists indicate datasets, brain regions, and sample sizes; group sizes are provided in Supplementary Table 1. Statistical thresholds are shown for each relevant step. Abbreviations: DE: differentially expressed, DEA: differential expression analysis, EC: entorhinal cortex, GWAS: genome-wide association study, miRNA: microRNA, mRNA: messenger RNA, NFT: neurofibrillary tangle, *p*adj: adjusted p-value, PFC: prefrontal cortex; TF: transcription factor.

Advanced AD pathology was associated with widespread alterations in cortical miRNA expression. We identified sixteen miRNAs that remained significant after Bonferroni correction (**Table 1**, **Fig. 2a**, **Supplementary Table 2**). The strongest and most consistent changes involved coordinated downregulation of the miR-132/212 cluster, with hsa-miR-132-3p (Bonferroni-adjusted *p*-value (*p_adj_*_)_ = 1.30 x 10^-13^), hsa-miR-132-5p (*p_adj_* _=_ 1.77 x 10^-10^), hsa-miR-212-3p (*p_adj_* _=_ 2.18 x 10^-12^) and hsa-miR-212-5p (*p_adj_* _=_ 5.12 x 10^-11^), representing the most significant associations across cohorts. Several members of the miR-129 family also showed reduced expression, whereas hsa-miR-193a-5p was the only significantly upregulated miRNA. In contrast, fewer miRNAs reached Bonferroni significance in the intermediate pathology meta-analysis. Across 1,048 individuals, four miRNAs were significantly associated with intermediate pathology **(Table 1**, **Fig. 2b, Supplementary Table 3)**: hsa-miR-125a-5p (*p_adj_* _=_ 7.73 × 10⁻⁹), hsa-miR-10a-5p (*p_adj_* _=_ 9.02 × 10⁻⁷), hsa-miR-132-3p (*p_adj_* _=_ 2.20 × 10⁻³), and hsa-miR-153-3p (*p_adj_* _=_ 0.014). Of these, hsa-miR-132-3p was Bonferroni significant in both the intermediate and advanced pathology analyses, with a larger effect size in advanced pathology (log₂FC = -0.75) than at the intermediate stage (log₂FC = -0.21). Notably, the distinction between intermediate- and advanced-stage associations was not absolute. Several miRNAs reaching Bonferroni significance only in advanced pathology already showed directionally concordant, nominally significant associations at the intermediate stage, although with generally smaller effect sizes. Conversely, hsa-miR-125a-5p, hsa-miR-10a-5p, and hsa-miR-153-3p showed their strongest associations with intermediate pathology but little or no evidence of differential expression in the advanced pathology analysis. Together, these findings suggest that miRNA dysregulation across NFT progression is stage-dependent but overlapping, with some changes detectable at intermediate stages and becoming more pronounced with advanced pathology, whereas others may be more prominent earlier in the pathological process.

**Fig 2:**
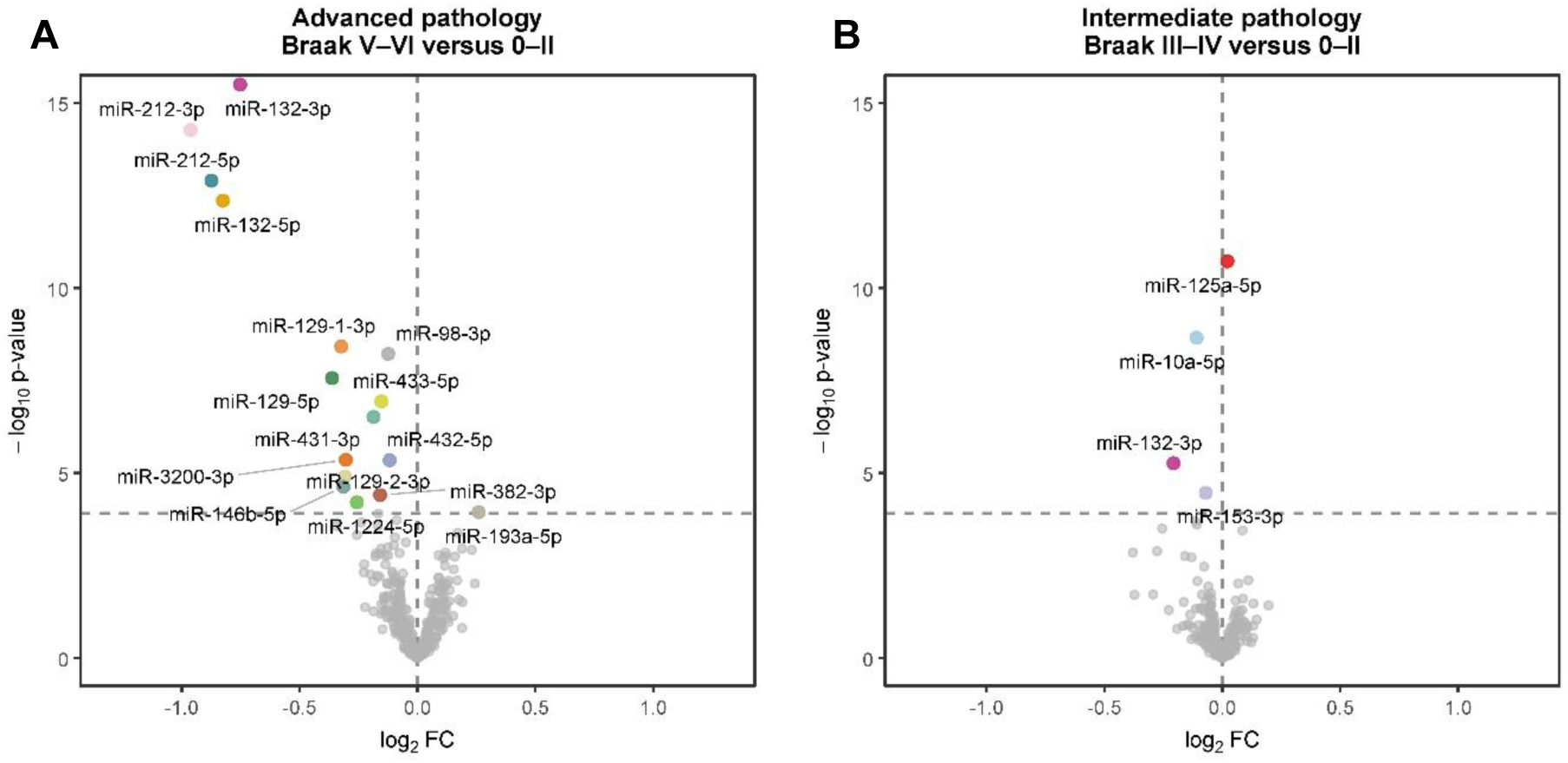
Meta-analysis identifies 19 miRNAs associated with AD cortical neurofibrillary tangle pathology. **a** Volcano plot showing differential miRNA expression between advanced Braak stage (V-VI, n = 690) and low Braak stage (0-II, n = 488) across seven datasets (N = 1,178). Colored points indicate the 16 miRNAs reaching Bonferroni significance. **b** Volcano plot showing differential miRNA expression between intermediate Braak stage (III-IV, n = 560) and low Braak stage (0-II, n = 488) across five datasets (N = 1,048). Colored points indicate the four Bonferroni significant miRNAs. X-axes show log2 fold change (log2 FC) and the y-axes show -log10 (*p*-value).

**Table 1:** Bonferroni-significant miRNAs associated with advanced and intermediate Braak stage. Differential expression analysis (DEA) results from random-effects meta-analysis for advanced (Braak V-VI versus 0-II) and intermediate (Braak III-IV vs. 0-II) pathology across cortical cohorts. Only miRNAs reaching Bonferroni significance in at least one comparison are shown, with Bonferroni significant findings shown in bold. Log2 fold change (log2FC), nominal *p*-values and Bonferroni-adjusted *p*-values (*padj*) are reported for both comparisons. Complete meta-analysis results for all 405 tested miRNAs are provided in Supplementary Tables 2 (advanced pathology) and 3 (intermediate pathology).

|  | miRNA ID | Advanced pathology meta-analysis |  |  | Intermediate pathology meta-analysis |  |  |
|---|---|---|---|---|---|---|---|
|  |  | log<sub>2</sub>FC | *p*-value | *p<sub>adj.</sub>* | log<sub>2</sub>FC | *p*-value | *p<sub>adj.</sub>* |
| **Common** | hsa-miR-132-3p | **-0.75** | **3.20x10<sup>-16</sup>** | **1.30x10<sup>-13</sup>** | **-0.21** | **5.44x10<sup>-6</sup>** | **2.20x10<sup>-3</sup>** |
| **Advanced Braak stage specific** | hsa-miR-212-3p | **-0.96** | **5.37x10<sup>-15</sup>** | **2.18x10<sup>-12</sup>** | -0.38 | 1.41x10<sup>-3</sup> | 0.569 |
|  | hsa-miR-212-5p | **-0.87** | **1.27x10<sup>-13</sup>** | **5.12x10<sup>-11</sup>** | -0.37 | 0.020 | 1.00 |
|  | hsa-miR-132-5p | **-0.83** | **4.38x10<sup>-13</sup>** | **1.77x10<sup>-10</sup>** | -0.25 | 3.17x10<sup>-4</sup> | 0.128 |
|  | hsa-miR-129-1-3p | **-0.32** | **3.82x10<sup>-9</sup>** | **1.55x10<sup>-6</sup>** | -0.11 | 8.31x10<sup>-3</sup> | 1.00 |
|  | hsa-miR-98-3p | **-0.12** | **6.05x10<sup>-9</sup>** | **2.45x10<sup>-6</sup>** | -0.11 | 2.46x10<sup>-4</sup> | 0.100 |
|  | hsa-miR-129-5p | **-0.36** | **2.73x10<sup>-8</sup>** | **1.10x10<sup>-5</sup>** | -0.05 | 0.095 | 1.00 |
|  | hsa-miR-433-5p | **-0.15** | **1.17x10<sup>-7</sup>** | **4.72x10<sup>-5</sup>** | -0.11 | 2.00x10<sup>-4</sup> | 0.081 |
|  | hsa-miR-431-3p | **-0.19** | **3.05x10<sup>-7</sup>** | **1.24x10<sup>-4</sup>** | -0.05 | 0.144 | 1.00 |
|  | hsa-miR-3200-3p | **-0.3** | **4.40x10<sup>-6</sup>** | **1.78x10<sup>-3</sup>** | -0.13 | 1.89x10<sup>-3</sup> | 0.766 |
|  | hsa-miR-432-5p | **-0.12** | **4.51x10<sup>-6</sup>** | **1.83x10<sup>-3</sup>** | -0.05 | 0.120 | 1.00 |
|  | hsa-miR-129-2-3p | **-0.31** | **1.24x10<sup>-5</sup>** | **5.02x10<sup>-3</sup>** | -0.04 | 0.066 | 1.00 |
|  | hsa-miR-146b-5p | **-0.31** | **2.37x10<sup>-5</sup>** | **9.58x10<sup>-3</sup>** | -0.03 | 0.093 | 1.00 |
|  | hsa-miR-382-3p | **-0.16** | **3.96x10<sup>-5</sup>** | **1.60x10<sup>-2</sup>** | -0.05 | 0.285 | 1.00 |
|  | hsa-miR-1224-5p | **-0.26** | **6.18x10<sup>-5</sup>** | **2.50x10<sup>-2</sup>** | -0.16 | 0.031 | 1.00 |
|  | hsa-miR-193a-5p | **0.26** | **1.14x10<sup>-4</sup>** | **4.62x10<sup>-2</sup>** | 0.10 | 0.165 | 1.00 |
| **Intermediate Braak stage specific** | hsa-miR-125a-5p | 0.01 | 0.842 | 1.00 | **0.02** | **1.91x10<sup>-11</sup>** | **7.73x10<sup>-9</sup>** |
|  | hsa-miR-10a-5p | 0.03 | 0.675 | 1.00 | **-0.11** | **2.23x10<sup>-9</sup>** | **9.02x10<sup>-7</sup>** |
|  | hsa-miR-153-3p | 0.00 | 0.922 | 1.00 | **-0.07** | **3.44x10<sup>-5</sup>** | **1.39x10<sup>-2</sup>** |

### Multi-omic integration identifies downstream transcriptional and proteomic targets of AD-associated miRNAs

To investigate the downstream consequences of AD-associated miRNA dysregulation, we separately integrated miRNAs significantly associated with advanced (n = 16) and intermediate (n = 4) pathology with matched mRNA (five cohorts) and proteomic (one cohort) datasets. Candidate targets were retrieved using the multiMiR database and retained only when the corresponding mRNA or protein were differentially expressed at Bonferroni and nominal significance levels, respectively, and there was a false discovery rate (FDR)-significant association between the expression levels of the miRNA and the corresponding mRNA or protein in the opposite direction to its upstream miRNA. This filtering strategy was consistent with the canonical repressive mode of miRNA action, whereby increased miRNA expression is expected to reduce target expression.

At the transcriptomic level, this approach identified 285 directionally consistent miRNA-mRNA interactions involving 13 miRNAs and 138 target mRNAs (**Supplementary Table 4**, **Fig 3**). Multiple targets were shared across the miR-132/212 family, including genes involved in synaptic organization, intracellular signaling and vesicle trafficking. Notable examples included *CLMN* and *TRIP10*, which showed strong inverse associations with hsa-miR-132-3p. In contrast, intermediate pathology yielded only a single significant regulatory interaction, linking hsa-miR-132-3p to the vesicle-trafficking gene *SYTL4* (**Supplementary Table 5**).

**Fig 3:**
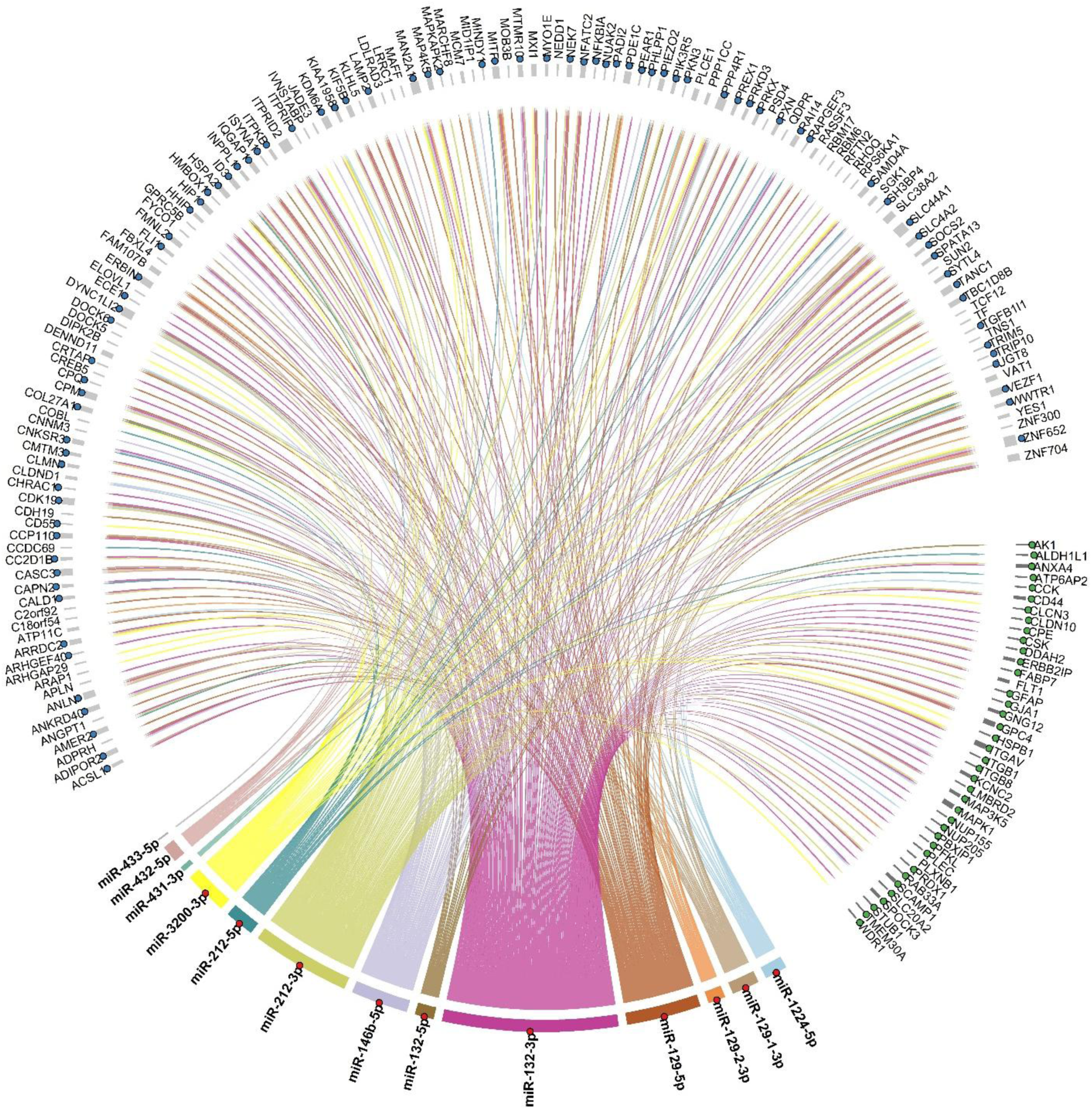
Integrated miRNA-target interaction network. Chord diagram showing significant miRNA-mRNA and miRNA-protein interactions for miRNAs associated with advanced tau pathology (Braak V-VI versus 0-II). Chords connect Bonferroni-significant miRNAs with downstream mRNA and protein targets identified through multi-omics integration. mRNA targets were required to be differentially expressed (*padj* < 0.05), show the expected inverse direction of effect, and exhibit a significant miRNA-mRNA association (*padj* < 0.05) across five matched RNA-sequencing cohorts. Protein targets were required to be differentially expressed (nominal p < 0.05), show the expected inverse direction of effect, and exhibit a significant miRNA-protein association (*padj* < 0.05) in ROSMAP. Chords are colored according to the corresponding miRNA. Colored circles adjacent to node labels indicate mediation at the miRNA (red), mRNA (blue), and protein (green) levels.

Integration with cortical proteomic data identified 55 significant miRNA-protein interactions involving 10 miRNAs and 40 proteins for advanced pathology (**Supplementary Table 6**, **Fig 3**), whereas only 29 interactions involving three miRNAs were detected during intermediate pathology (**Supplementary Table 7**). Consistent with the transcriptomic analyses, the majority of proteomic associations involved members of the miR-132/212 family, including proteins implicated in MAPK signaling and synaptic function.

### miRNA dysregulation mediates downstream molecular changes associated with AD pathology

To determine whether changes in gene and protein expression associated with pathology were mediated by disease-associated miRNAs, we performed causal mediation analyses for all significant miRNA-target pairs from the advanced pathology analysis in the four PFC cohorts. Among the 285 miRNA–mRNA interactions we had previously identified, 122 interactions involving six miRNAs (hsa-miR-132-3p, hsa-miR-132-5p, hsa-miR-212-3p, hsa-miR-212-5p, hsa-miR-146b-5p, hsa-miR-3200-3p) and 84 target mRNAs met the criteria for replicated, predominantly miRNA-mediated associations (**Supplementary Table 8**, **Fig. 3**). These interactions showed significant average causal mediation effects (ACME) with unadjusted *p* < 0.05 in at least three out of four tested cohorts, with the same direction. Representative examples included regulatory axes linking hsa-miR-132-3p to *CLMN* and hsa-miR-212-3p to *MAP4K5*.

At the proteomic level, 26 of the 55 miRNA–protein interactions showed a significant predominantly miRNA-mediated pattern in the ROSMAP cohort (**Supplementary Table 9**, **Fig 3**). These interactions involved six miRNAs (hsa-miR-132-3p, hsa-miR-212-3p, hsa-miR-212-5p, hsa-miR-146b-5p, hsa-miR-3200-3p, and hsa-miR-1224-5p), of which five miRNAs also featured in our miRNA-mRNA mediation analysis.

When we examined the miRNA-target interactions with intermediate pathology, we saw no evidence of mediation for the single miRNA-mRNA association we had identified between hsa-miR-132-3p and *SYTL4*, nor for the 29 miRNA-protein relationships after multiple-testing correction.

### Consensus co-expression network analysis identifies transcriptional programs associated with AD-related miRNAs

To identify coordinated transcriptional programs associated with AD-related miRNAs, we next performed consensus weighted gene co-expression network analysis (WGCNA) across the four postmortem prefrontal cortex (PFC) transcriptomic datasets. Following residualization for technical and demographic covariates and variance filtering, 3,180 genes shared across all cohorts were used to construct a consensus co-expression network, resulting in eleven common co-expression modules across the four cohorts.

Correlation of module eigengenes with Braak stage and the expression levels of the 16 advanced pathology-associated miRNAs identified highly reproducible module-trait relationships across cohorts (**Fig. 4a**, **Supplementary Fig. 1**). Braak stage showed positive correlations with the green, blue and yellow modules and negative correlations with the turquoise and black modules. In contrast, the miR-132/212 family, which we had previously shown were downregulated in advanced pathology, displayed the opposite pattern, exhibiting strong negative correlations with the green and blue modules, as well as the red module, and positive correlations with the turquoise and black modules, as well as the purple module. Similar correlation patterns were observed for several additional advanced pathology-associated miRNAs, including miR-146b-5p, miR-1224-5p and miR-433-5p, indicating that these miRNAs are associated with coordinated but distinct transcriptional programs during AD progression.

**Fig. 4:**
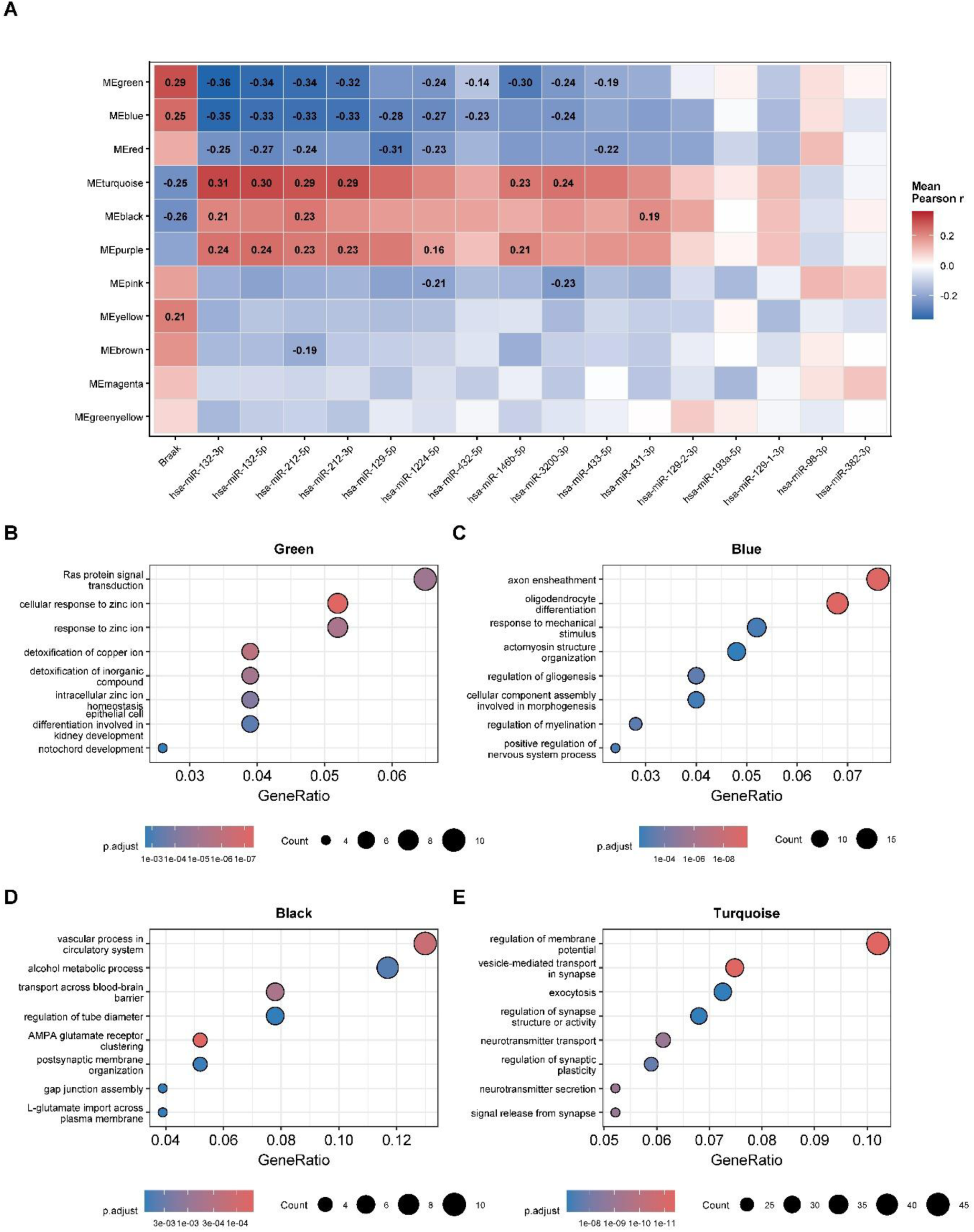
Consensus gene co-expression network analysis identifies transcriptional programs associated with AD-related miRNAs. **a** Heatmap showing correlations between consensus module eigengenes, Braak stage, and prioritized miRNAs from the advanced pathology analysis across four independent postmortem prefrontal cortex transcriptomic datasets (BDR-1, PITT-ADRC, UKBBN, and ROSMAP). Colors represent the mean Pearson correlation coefficient across cohorts, with red representing positive correlations and blue representing negative correlations. Numerical values indicate reproducible associations with nominal *p* < 0.05 with concordant directions of effect in at least three of the four cohorts. **b-e** Gene Ontology (GO) over-representation analysis of genes within the **b** green, **c** blue, **d** black, and **e** turquoise consensus co-expression modules. Dot size represents the number of genes annotated to each GO term, color indicates the adjusted *p*-value (Benjamini-Hochberg), and the x-axis represents the gene ratio.

Gene Ontology (GO) enrichment analysis demonstrated clear functional differences between the four modules associated with both Braak stage and miRNA expression (**Fig. 4b-e**, **Supplementary Table 10**). The turquoise module, positively correlated with the miR-132/212 family and negatively correlated with Braak stage, was particularly enriched for neuronal and synaptic processes, including regulation of membrane potential, vesicle-mediated transport in synapses, exocytosis, neurotransmitter transport and regulation of synaptic plasticity (**Fig. 4e**). Conversely, the blue module, which showed the opposite correlation pattern, was most enriched for oligodendrocyte differentiation, axon ensheathment, regulation of myelination and gliogenesis (**Fig. 4c**). The black module was enriched for vascular and blood-brain barrier processes, including vascular function, regulation of tube diameter and transport across the blood-brain barrier (**Fig. 4d**), whereas the green module was enriched for metal ion homeostasis and detoxification pathways, including cellular responses to zinc and copper ions and intracellular zinc homeostasis (**Fig. 4b**).

### miRNA-mediated mRNA targets converge on NFT-associated glial transcriptional signatures

Next, we took the 122 mRNA targets that had shown NFT-associated expression changes in bulk tissue and were predominantly mediated by NFT-associated miRNA expression and examined their expression profiles in relation to NFT pathology across 49 human prefrontal cortex cell clusters reported by Mathys *et al*.^11^. Significant enrichment was restricted to astrocytes, oligodendrocytes, and microglia, with astrocytes showing the strongest evidence (**Fig. 5**, **Supplementary Table 11**). In astrocytes (Ast), 36 of the 50 targets represented in the cluster-specific gene background overlapped with NFT-associated differentially expressed genes (OR = 4.72; *p*_adj =_ 6.86 × 10⁻⁶). These included *NFKBIA, HSPA2, MITF, INPPL1, LAMP2, ECE1*, and *ITPKB*.

**Fig 5:**
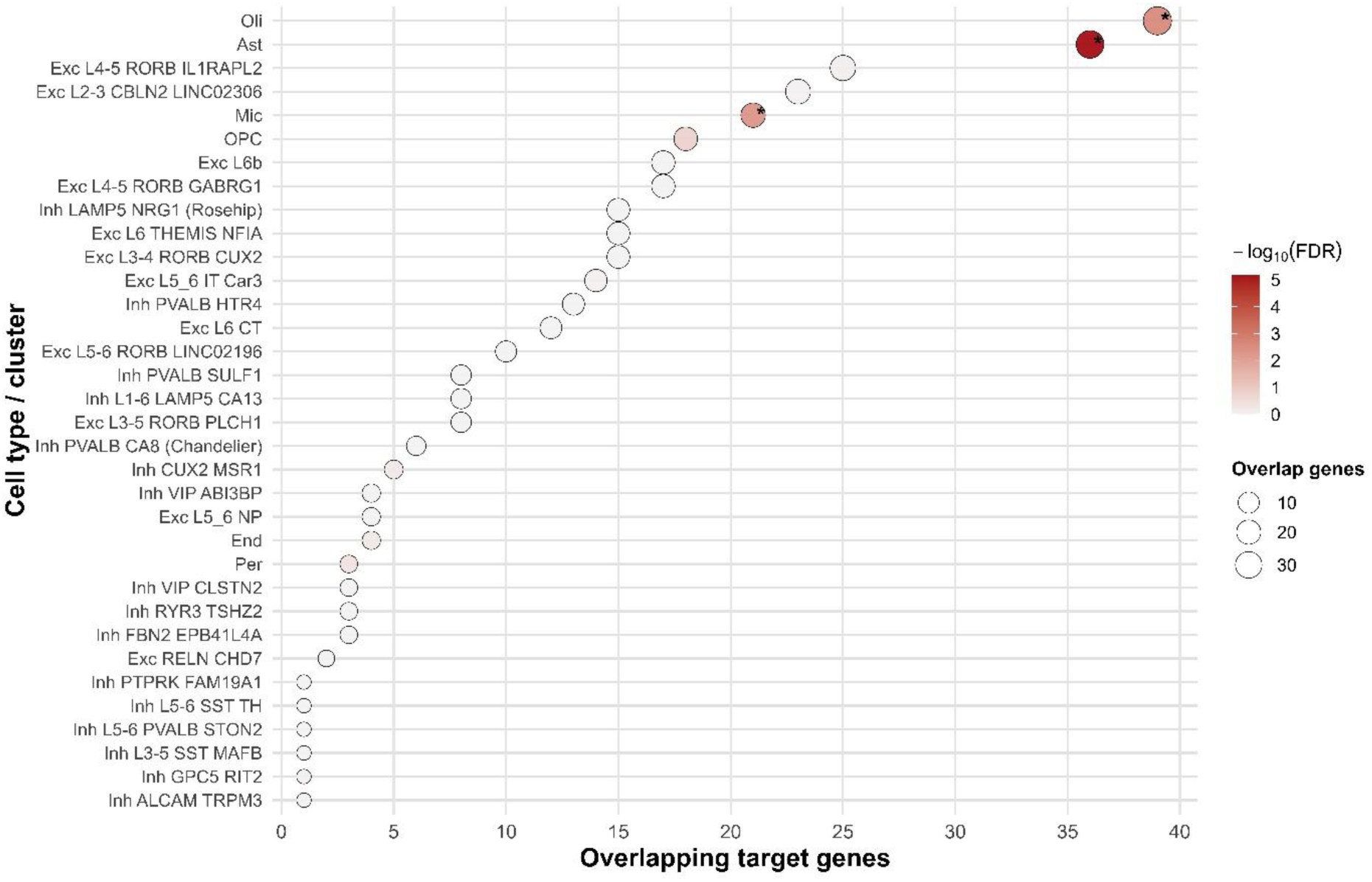
Cell-type enrichment of AD-associated miRNA target genes. Dot plot showing enrichment of inversely associated AD miRNA target genes among differentially expressed NFT-associated genes across single-cell transcriptomic cell clusters. Dot size represents the number of overlapping genes, and color indicates enrichment significance (−log₁₀ *p*adj from one-sided hypergeometric tests). Asterisks denote significant enrichment (*p*adj < 0.1).

Significant enrichment was also observed in mature oligodendrocytes (Oli; 39 of 60 targets; OR = 2.74; *p*_adj =_ 2.63 × 10⁻³) and microglia (Mic; 21 of 57 targets; OR = 2.83; *p*_adj =_ 4.92 × 10⁻³). In oligodendrocyte precursor cells (OPC), 18 of 48 targets overlapped with NFT-associated differentially expressed genes, but enrichment did not survive multiple-testing correction (OR = 2.07; *p*_adj =_ 0.163).

No neuronal cluster showed significant enrichment after multiple-testing correction. The largest overlap among excitatory populations occurred in Exc L4-5 RORB IL1RAPL2, involving 25 of 39 targets (64.1%), whilst among inhibitory populations Inh LAMP5 NRG1 (Rosehip) showed the largest overlap, involving 15 of 35 targets (42.9%) **(Fig. 5**, **Supplementary Table 11)**. These overlaps primarily involved targets linked to miR-132-3p and miR-212-3p, corresponding to 21 and 13 mRNAs, respectively, in the excitatory cluster, and 14 and seven genes in the inhibitory cluster, respectively, with some genes linked to both miRNAs.

The two neuronal overlap sets shared 10 targets, with 15 additional targets found only in the excitatory set and five only in the inhibitory set. Targets exclusive to the excitatory set included *DYNC1LI2, KIF5B, MAPKAPK2, MTMR10*, and *ISYNA1*, whereas those exclusive to the inhibitory set were *ANLN, CPQ, ECE1, FMNL2*, and *ZNF652*. miR-132-5p was represented only in the excitatory overlap set, through *ISYNA1*.

### Multi-omic network integration identifies cell-type-specific regulatory architectures and recurrent miRNA hubs

To characterize AD-associated regulatory shifts, we integrated single-nucleus transcriptomics, transcription factor (TF)-miRNA relationships, and cortical mediation models to reconstruct directed TF-miRNA-mRNA networks across 35 brain cell types. The global architecture comprised 1,562 localized nodes and 2,812 directed edges (**Supplementary Table 12**). Enforcing directional constraints revealed a heavily repressive global topology (78.8% of interactions), cascading into canonical post-transcriptional miRNA-target decay (**Supplementary Table 13**). Network connectivity varied substantially; the most expansive, hyper-connected sub-networks localized to excitatory neurons (*e*.*g*., Exc L2-3 CBLN2 LINC02306; 123 nodes, 250 edges) and macroglia (astrocytes and oligodendrocytes; >100 nodes, >200 edges) (**Supplementary Table 12**).

Topological centrality analyses identified specific miRNAs as the strong master regulators connecting upstream transcriptional shifts to downstream mRNA repression (**Supplementary Table 14**). The miR-132/212 cluster emerged as a recurrent hub across vulnerable lineages. Specifically, miR-132 exhibited peak topological dominance in deep-layer excitatory neurons (Exc L4-5 RORB IL1RAPL2; total degree = 75) and oligodendrocytes (total degree = 71). Similarly, miR-146b operated as a secondary master node, driving dense target subnetworks in upper-(total degree = 73) and deep-layer (total degree = 69) excitatory populations.

The pattern of regulatory edge similarity suggests that cell identity constrains miRNA-associated network architecture (**Fig. 6**, **Supplementary Table 15**, **Supplementary Table 16**). Closely related excitatory neuronal subtypes shared the greatest proportion of regulatory interactions, whereas inhibitory populations were more heterogeneous, whilst glial and vascular populations showed more distinct architectures. The consistently lower edge than node similarity indicates that cell types often share regulatory components but differ in how these components are connected. Thus, cell-type specificity may arise through context-dependent wiring of a partially shared regulatory repertoire rather than through entirely distinct sets of regulators. These networks, however, represent inferred cell-type-contextualized architectures rather than direct measurements of cell-type-specific miRNA activity. Finally, cross-referencing downstream targets against CellChatDB revealed negligible overlap, identifying only ADIPOR2 as a receptor (for the ADIPOQ ligand). This indicates the reconstructed regulatory networks predominantly capture intracellular molecular dynamics rather than extracellular signaling events.

**Fig 6:**
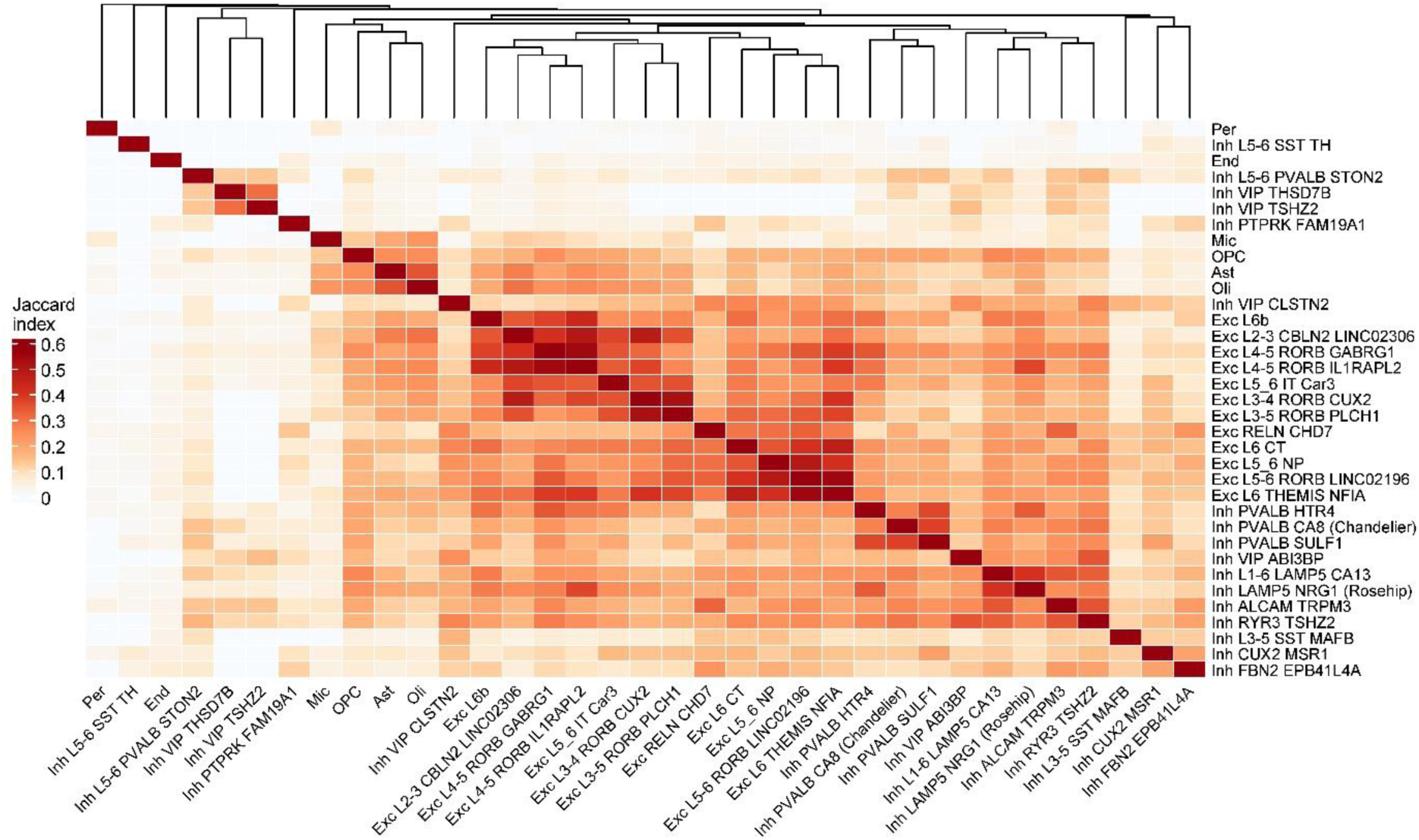
Regulatory edge similarity across cell-type-specific networks. Heatmap showing pairwise Jaccard similarity of directed regulatory edges across the 35 reconstructed cell-type-specific transcription factor (TF)-miRNA-mRNA networks. Higher Jaccard values indicate greater overlap in regulatory interactions between cell types. Cell types were hierarchically clustered using average linkage on Jaccard distance (1 − Jaccard similarity), with the resulting dendrogram shown above the heatmap.

### Cis-miQTL mapping identifies genetic regulators of Braak-associated miRNAs, but no evidence of co-localization with AD risk loci

To investigate whether the expression of Braak-associated miRNAs was under genetic regulation, we performed cis-miRNA quantitative trait locus (miQTL) mapping for the 16 miRNAs associated with advanced pathology using matched genotype and miRNA expression data from BDR-2, UKBBN, ROSMAP, and PITT-ADRC (total n = 1,092). Following inverse-variance meta-analysis of 62,373 cis-miQTLs present in at least three of the four cohorts, 31 were associated with seven of the advanced pathology-associated miRNAs at an unadjusted *p* < 1 × 10^-3^ (**Supplementary Table 17**).

We next tested whether these cis-miQTL signals shared causal variants with established AD susceptibility loci from the consensus meta-analysis of genome-wide association studies for AD^12^ . Bayesian colocalization analysis revealed that none of these miQTL loci showed strong evidence of colocalization (PP4 > 0.9) with the nominated AD genomic regions, providing no evidence that cis-genetic regulation of these miRNAs shares causal signals with established AD risk loci.

## Discussion

This study provides a large-scale, multi-omic assessment of cortical miRNA dysregulation in association with NFT pathology in AD. Rather than identifying a uniform miRNA signature that simply increases with disease severity, our findings point to a more dynamic regulatory landscape in which some miRNA alterations are associated with particular pathological stages, while others persist across the disease continuum. Within this landscape, the miR-132/212 cluster emerged as the most consistent signal and repeatedly converged with downstream transcriptional, proteomic and cell-type-associated changes. These observations suggest that miRNA dysregulation is embedded within broader molecular responses to NFT pathology and may contribute to their organization across different cellular contexts.

The prominent reduction of miR-132/212 strongly corroborates previous evidence from human AD brain studies. Lau and colleagues identified miR-132-3p as one of the most strongly altered miRNAs in AD in both hippocampal and prefrontal cortical tissue and showed that its reduction was already evident at Braak stages III-IV, before widespread loss of neuron-associated miRNAs^13^. Wong and colleagues independently demonstrated downregulation of both miR-132 and miR-212 in AD temporal cortex and CA1 neurons and linked loss of the cluster experimentally to neuronal vulnerability through PTEN-FOXO3a-related signalling^14^. Together, these studies establish miRNA dysregulation as a reproducible feature of the AD brain and provide strong external support for the prominent miR-132/212 signal observed here.

Our comparison of intermediate and advanced pathology adds an important dimension to this literature. miR-132-3p was unique in being significantly associated with both pathological stages, whereas several miRNAs were confined to either intermediate or advanced pathology. This pattern argues against a simple monotonic model of miRNA dysregulation during AD progression. Instead, distinct post-transcriptional responses may characterize different pathological states. The persistence of miR-132-3p is consistent with the earlier observations of Lau *et al*. and suggests that disruption of the miR-132 pathway may arise relatively early and remain detectable as NFT pathology becomes more widespread. Conversely, intermediate-stage-specific signals may represent transient responses, compensatory mechanisms, or molecular states that are subsequently obscured by later pathological processes. Because NFT Braak-stage comparisons are cross-sectional, however, these patterns should not be interpreted as longitudinal trajectories within individuals. This distinction is particularly important when considering miRNAs that appear altered at intermediate but not advanced stages.

There is substantial experimental evidence supporting the biological relevance of miR-132 loss to AD-related processes. miR-132 is an activity-regulated neuronal miRNA involved in dendritic maturation, spine formation and neuronal integration^15^. In AD models, loss of miR-132 derepresses ITPKB and exacerbates both amyloid and tau pathology through downstream ERK1/2 and BACE1-related mechanisms^7^, while deficiency of the miR-132/212 cluster increases Aβ production and plaque deposition^16^. Conversely, restoration of miR-132 has been shown to reduce pathological forms of tau and protect neurons in experimental tauopathy models^17^, and to rescue impaired adult hippocampal neurogenesis and memory phenotypes in AD models^18^. Our findings therefore extend a well-established experimental literature by showing that miR-132/212 dysregulation is reproducible across multiple human cortical cohorts and sits within a much broader network of disease-associated molecular changes.

Our multi-omic approach enabled us to link AD-associated miRNA dysregulation to downstream changes in transcript and protein expression. Rather than relying solely on predicted target relationships, we prioritized interactions that were supported by disease-associated changes in the target transcript and/or protein and that showed the inverse direction expected for canonical miRNA regulation. This identified a broad downstream regulatory footprint, particularly for the miR-132/212 family. The integration of protein abundance is especially valuable because transcript and protein changes in AD are only partially concordant. Large-scale human brain proteomic studies have demonstrated that major AD-associated protein networks can be poorly represented at the RNA level, emphasizing that transcriptomic and proteomic analyses capture complementary aspects of disease biology^19^. Thus, the recurrence of miRNA-associated effects across both RNA and protein levels provides stronger support for their biological relevance than transcript-level associations alone.

Our mediation analyses provide further evidence that miRNA variation may participate in these downstream molecular changes. A substantial proportion of the prioritized transcript and protein relationships were consistent with statistical mediation through miRNA expression. We interpret this as supporting a regulatory model in which miRNA dysregulation contributes to the relationship between NFT pathology and downstream molecular abundance, rather than as proof of causal direction. Given that our mediation analyses were performed cross-sectionally, we also cannot exclude feedback regulation, common upstream drivers or changes in cellular composition. Nevertheless, when statistical mediation converges with inverse expression relationships, replicated human associations and experimental evidence for miRNA function, it provides a useful framework for prioritizing candidate regulatory axes for mechanistic investigation.

The opposing associations of miR-132/212 and Braak pathology with neuronal/synaptic and oligodendrocyte/myelination-related co-expression programs suggest that miRNA dysregulation accompanies changes across cortical cellular programs. The neuronal component is consistent with the established role of miR-132 in activity-dependent maturation and synaptic plasticity, while the oligodendrocyte component aligns with human single-nucleus studies identifying AD-associated regulatory modules linked to *APOE* and *CLU* ^20^ and subtype-specific alterations in astrocytes and oligodendrocytes^21^.

Integration with NFT-associated single-nucleus profiles reinforced this glial component, with significant enrichment observed in astrocytes, oligodendrocytes, OPCs, and microglia, and most strongly in astrocytes. miR-132-3p and miR-212-3p had the broadest target representation. The glial overlaps included regulators of lysosomal function and autophagy (*LAMP2*), protein quality control (*HSPA2*), inositol signaling (*INPPL1*, *ITPKB*), and inflammatory signaling (*NFKBIA*). Additional connections included miR-146b-5p-linked *DOCK5* in Rac signaling and miR-132-5p-linked *UGT8* in oligodendrocyte myelin lipid synthesis. These functional links complement the heterogeneous glial responses previously described in AD cortical studies^11,22^.

Although neuronal enrichment did not survive multiple-testing correction, Exc L4–5 RORB IL1RAPL2 and Inh LAMP5 NRG1 showed substantial target overlaps, predominantly linked to miR-132-3p and miR-212-3p. Shared targets included *HIP1*, *PXN*, and *CAPN2*, connecting these descriptive overlaps to endocytosis, cell adhesion, and calcium-dependent proteolysis. Distinct targets included the intracellular transport genes *DYNC1LI2* and *KIF5B* in the excitatory overlap set and the actin regulator *FMNL2* in the inhibitory set. These findings identify candidate neuronal contexts for further investigation.

These analyses infer cellular context from bulk cortical miRNA measurements, single-nucleus mRNA profiles, and cell-type-specific TF information; they do not directly demonstrate mature miRNA dysregulation within individual cell populations. Conventional single-cell and single-nucleus RNA-seq poorly capture mature miRNAs. Elias *et al*. developed primiReference to quantify primary miRNA transcripts and identified cell-type-specific changes in 5xFAD mouse cortex^23^ , but precursor transcription is not equivalent to mature miRNA abundance or activity. Direct spatial or cell-type-resolved mature miRNA profiling will therefore be needed to establish the cellular origin of these signals.

The pattern of regulatory edge similarity suggests that cell identity constrains miRNA-associated network architecture. Closely related excitatory neuronal subtypes shared the greatest proportion of regulatory interactions, whereas inhibitory populations were more heterogeneous and glial and vascular populations showed more distinct architectures. The consistently lower edge than node similarity indicates that cell types often share regulatory components but differ in how these components are connected. Thus, cell-type specificity may arise through context-dependent wiring of a partially shared regulatory repertoire rather than through entirely distinct sets of regulators. These networks, however, represent inferred cell-type-contextualized architectures rather than direct measurements of cell-type-specific miRNA activity. This interpretation aligns with increasingly detailed human single-cell atlases showing that pathological responses can differ substantially even between closely related neuronal subtypes and that selective cellular vulnerability is strongly dependent on transcriptional identity^11^.

Our study has several strengths, most notably the large sample size, utilizing seven independent datasets. In addition, our multi-omic approach has allowed us to follow prioritized miRNAs across transcriptional, proteomic, mediation and cell-type-associated regulatory analyses. The repeated prioritization of miR-132/212 by these analytically distinct approaches is therefore more informative than its prominence in any individual analysis. Several limitations should nevertheless be considered. First, the study is based on postmortem tissue and cannot establish the temporal sequence of molecular changes within individuals. Pathological stage provides a biologically meaningful ordering of NFT burden but is not equivalent to a longitudinal disease trajectory. Second, bulk miRNA measurements are influenced by cellular composition, and the downstream single-nucleus analyses provide cell-type context rather than direct evidence of cell-type-specific mature miRNA dysregulation. Although we have used surrogate variables to indirectly account for cell proportion changes between samples, we are not able to assess miRNAs (and their target mRNA/protein targets) in specific cell populations. Third, protein-level relationships were evaluated in only one cohort and lack the same degree of cross-cohort replication available for the RNA analyses. Fourth, network reconstruction depends on existing TF-miRNA and miRNA-target annotations, which are incomplete and may not capture context-specific regulatory activity. Finally, neither statistical mediation nor network centrality establishes true causal molecular regulation. Experimental perturbation of prioritized miRNA-target relationships in relevant neuronal and glial systems will be required to distinguish mechanisms that actively contribute to disease progression from molecular responses to accumulating pathology.

Overall, our findings support a model in which cortical miRNA dysregulation in AD is both pathology-stage- and cellular-context-dependent. miR-132/212 represents the strongest point of convergence between the present human data and previous mechanistic studies, while the emergence of miR-146b, the substantial glial target enrichment and the cell-type-dependent regulatory networks broaden the landscape beyond the established neuronal pathway. Rather than pointing to a single miRNA-mediated mechanism, our results suggest that a relatively restricted set of dysregulated miRNAs may influence distributed molecular programs whose consequences depend on cellular context. Direct spatial and cell-type-resolved profiling of mature miRNAs, combined with perturbation of the prioritized regulatory axes, will be important for establishing which components of this network are causal contributors to AD pathology and which reflect adaptive or secondary responses.

## Online methods

### Brain banks and cohort characteristics

Postmortem brain tissue from the entorhinal cortex (EC) or the prefrontal cortex (PFC) was utilized from six cohorts: the MRC London Neurodegenerative Diseases Brain Bank (LBBND), the UK Brain Bank Network (UKBBN), the Brains for Dementia Research (BDR) program, the Pittsburgh Alzheimer’s Disease Research Center (PITT-ADRC), the Oxford Project to Investigate Memory and Ageing (OPTIMA), and the Religious Orders Study and Memory and Aging Project (ROSMAP). For LBBND, UKBBN, BDR, PITT-ADRC, and OPTIMA, tissue collection and use were approved by the respective institutional review boards. For the ROSMAP cohort, previously generated omics data were obtained from the AMP-AD Knowledge Portal (https://adknowledgeportal.synapse.org) under approved data use agreements. Across all six cohorts, donors underwent standardized neuropathological assessments, and all samples were processed and stored according to cohort-specific protocols. Demographic and neuropathological characteristics are provided in **Supplementary Table 1.**

### Neuropathological outcomes and pathology staging

The primary neuropathological outcome examined was the progression of neurofibrillary tangle (NFT) burden composed of hyperphosphorylated tau, pathologically evaluated across postmortem brain tissues using the six-tier Braak NFT staging system. To capture distinct clinical and pathological phases of neurodegeneration, Braak staging scores were grouped into three progressive categories: low pathology (stages 0-II), intermediate pathology (stages III-IV), and advanced pathology (stages V-VI). Downstream differential expression models evaluated two explicit pathological contrasts, specifically testing intermediate versus low pathology to capture early-to-moderate changes, and advanced versus low pathology to identify late-stage alterations.

Standardized Braak staging records were harmonized and available across all six sample cohorts. To ensure that molecular associations with NFT progression were not driven by co-occurring neurodegenerative or cerebrovascular conditions, comprehensive sensitivity analyses were executed. These validation models systematically excluded donors displaying substantial comorbid features, including significant vascular pathology, Lewy body pathology, or transactive response DNA-binding protein 43 (TDP-43) inclusions, where such deep neuropathological metadata were available within the respective cohort repositories.

### Small RNA sequencing

For the BDR cohort, laboratory experiments and downstream analyses were performed independently in two batches (BDR-1: N = 81; BDR-2: N = 420), which were therefore treated as separate datasets, resulting in seven datasets across six cohorts. For LBBND, BDR-1, and BDR-2 cohorts, total RNA was isolated from approximately 30 mg of dissected brain tissue using QIAzol Lysis Reagent and the miRNeasy Mini Kit (Qiagen), according to the manufacturer’s instructions. RNA integrity was assessed using the Agilent RNA ScreenTape assay on an Agilent 4200 TapeStation system. RNA extraction and processing procedures for the PITT-ADRC, UKBBN, OPTIMA, and ROSMAP cohorts have been described previously^24–27^.

Small RNA libraries were subsequently prepared using the NEXTFLEX Small RNA-Seq Kit v3 (PerkinElmer) and subjected to next-generation sequencing. LBBND samples were sequenced on an Illumina HiSeq 2500, UKBBN, BDR-1, BDR-2, and PITT-ADRC samples on an Illumina NovaSeq 6000, and OPTIMA samples on an Illumina HiSeq 4000. Sequencing generated 50-bp single-end reads across all cohorts, except for BDR-1, for which paired-end sequencing was performed. For the ROSMAP cohort, RNA was extracted using TRIzol, and miRNA libraries were prepared using the NEBNext Multiplex Small RNA Library Prep Kit for Illumina (New England Biolabs) and sequenced on an Illumina HiSeq 3000, as previously described^27^.

Raw sequencing files were processed through the miRDeep2 software pipeline^28^ to identify and quantify microRNA (miRNA) species. The core mapping engine generated non-redundant sequence profiles per sample, which were aligned to annotated human mature and precursor miRNA sequences from miRBase. For miRNAs derived from genomic loci with multiple hairpin precursors, expression values were collapsed and aggregated using the *WGCNA* R package (v1.74)^29^. Low-abundance transcripts were filtered out using the function filterByExpr in the *edgeR* (v4.8.0) R package^30^ to exclude noise from downstream analyses; miRNAs were retained only if they maintained a baseline of at least five raw reads in a minimum of 75% of the total sample population. Following filtering, remaining counts were normalized for variable library size using the trimmed mean of M-values method and converted to log_2 c_ounts per million values for statistical modeling.

Prior to data modeling, clinical and phenotypic covariates were systematically harmonized across all contributing brain banks. Missing postmortem interval (PMI) data points were handled via cohort-specific median imputation. To account for non-linear artifacts, continuous variables for PMI and RNA integrity number (RIN) were stratified into quartile-based categorical tiers, while sex, self-reported ethnicity, source brain bank, and pathological Braak stage were encoded as discrete categorical factors. Confounding variance was assessed by evaluating pairwise correlations between the first ten principal components of the normalized expression matrix and all compiled clinical metadata. Variables displaying significant associations with major axes of expression variation were explicitly included as fixed effects in downstream cohort-specific differential expression models. Residual unmeasured variation and hidden batch effects were captured via surrogate variable analysis using the *sva* package (v3.58.0)^31^. Five latent surrogate variables were estimated and integrated into the final linear regression models to serve as statistical adjustments and reference-free proxies for cell-type composition.

### RNA sequencing

For the UKBBN, BDR-1, PITT-ADRC, OPTIMA, and ROSMAP cohorts, ribosomal RNA was systematically depleted from total RNA extracts using the Illumina Ribo-Zero Plus rRNA Depletion Kit. Stranded library preparation was subsequently completed using the Illumina RNA Prep Kit. Final library quality, concentration, and size distributions were rigorously evaluated via the Qubit 1X dsDNA High Sensitivity Assay and the Agilent D1000 ScreenTape system on a TapeStation platform. Next-generation sequencing was executed on the Illumina NovaSeq 6000 platform for the UKBBN, BDR-1, PITT-ADRC, and OPTIMA cohorts, and across standardized Illumina sequencing architectures for the ROSMAP repository.

Raw sequencing reads were subjected to initial quality control profiling using FastQC (v0.13.0). Automated trimming was subsequently deployed to strip sequencing adapters and eliminate low-quality terminal bases. The remaining high-quality reads were aligned to the human reference genome (hg38 assembly) using the splice-aware STAR aligner. To ensure analytical consistency with the small RNA profiling pipeline, raw expression values were filtered to exclude low-abundance transcripts, normalized for variable library depth using the trimmed mean of M-values method within the *edgeR* package (v4.8.0)^30^, and transformed to log_2 c_ounts per million values prior to downstream differential expression analysis.

### Proteomics

To complement the transcriptomic datasets, we utilized a subset of previously generated proteomic data from the dorsolateral PFC in the ROSMAP cohort (n = 400)^19^. In brief, high-throughput proteomic profiling was performed using tandem mass tag (TMT)-based quantitative mass spectrometry. Tryptic peptides from individual donor samples were labeled with isobaric TMT reagents, pooled into multiplexed channels, and analyzed via high-resolution liquid chromatography-tandem mass spectrometry. Raw reporter ion intensities were extracted, and systematic variations were resolved by normalizing values both within and across experimental multiplex batches. The resulting reporter intensities were log_2-_transformed and statistically adjusted for technical covariates and latent batch effects. This analytical workflow yielded a finalized matrix of relative protein abundances, tracking thousands of high-confidence quantified proteins across the ROSMAP donor population.

### Genotyping data

Genotyping was performed using the Illumina Global Screening Array (GSA) for the UKBBN, BDR-1, BDR-2 and PITT-ADRC samples. For ROSMAP, genotype data generated using the Affymetrix GeneChip 6.0 and Illumina HumanOmniExpress arrays were obtained from the Synapse repository.

Genotype quality control (QC) was performed both before and after imputation. Duplicate samples were identified using KING (Kinship-based INference for GWAS; v2.2.7)^32^, and for duplicate pairs, the sample with the higher genotype missingness (>10%) was excluded. Samples with >10% missing genotypes were subsequently removed using PLINK v1.9 (--mind 0.1)^32,33^. Multiallelic variants were excluded, retaining only biallelic variants. Variants with < 5% missing genotypes (--geno 0.05) or deviation from Hardy–Weinberg equilibrium (HWE; P < 1 × 10^-6^) were removed. Variants were additionally filtered based on minor allele frequency (MAF).

Individual heterozygosity was assessed using autosomal variants after filtering for MAF, HWE, and genotype missingness, followed by linkage disequilibrium (LD) pruning. Samples showing excessive heterozygosity, defined as an absolute inbreeding coefficient |F| > 0.2, were excluded. Relatedness was assessed using KING, with samples showing kinship coefficients > 0.1 designated for exclusion; no related samples exceeding this threshold were identified. Genetic ancestry was evaluated by merging the study genotypes with 1000 Genomes reference data followed by principal component analysis (PCA) using PLINK v2.0. All study samples were clustered with the European-ancestry reference population, and no ancestry outliers were identified.

Prior to imputation, variant positions and alleles were checked against European reference data from the 1000 Genomes Project using the HRC/1000G imputation preparation script (HRC-1000G-check-bim.pl) recommended by the Michigan Imputation Server.

Genotypes were imputed against the 1000 Genomes Phase 3 v5 reference panel. Genotype datasets were harmonized to the hg19 genome build before imputation. ROSMAP genotypes, originally mapped to hg18, were lifted over to hg19 before QC and imputation. Following imputation, variants with imputation quality R^2^ < 0.3 were removed from dosage VCF files using PLINK v2.0, and the genotype QC procedure was repeated. The resulting genotype data were subsequently lifted over from hg19 to hg38 using UCSC LiftOver and the corresponding chain file to match the genome build of the expression data.

### Data analysis

All analyses were performed in R (v 4.5.2) unless otherwise specified. Meta-analyses of miRNA and mRNA expression were considered primary analyses, with statistical significance defined as Bonferroni-adjusted *p* (*p*_adj)_ < 0.05. All other analyses were considered secondary, with significance thresholds defined, as appropriate, as *p*_adj <_ 0.05, and described in the analyses below. A Benjamini-Hochberg (BH) false discovery rate (FDR) threshold of 0.10 was used for the exploratory cell-type analyses.

### Differential expression analysis

To identify pathological molecular alterations associated with disease progression, differential expression analyses were conducted using the *limma* R package^34^ (v3.66.0) across the individual molecular tiers. Models were fitted separately for each data modality in each dataset, encompassing for miRNAs the UKBBN, LBBND, BDR-1, BDR-2, PITT-ADRC, OPTIMA, and ROSMAP datasets, for mRNAs the UKBBN, BDR-1, PITT-ADRC, OPTIMA, and ROSMAP datasets, and for proteins the ROSMAP dataset. All core linear regression models were adjusted for age at death, sex, and the five previously estimated latent surrogate variables to control for unmeasured sources of variation, including cell-type composition heterogeneity.

To account for cohort-specific technical variation, source brain bank and sequencing plate batch variables were explicitly integrated as covariates in instances where they displayed significant associations with the first ten principal components of the expression matrices. To map molecular signatures across distinct phases of tau pathology, analyses were stratified by NFT Braak stage, testing for significant differential expression of miRNAs, mRNA transcripts and proteins in intermediate-stage disease (Braak stages III-IV versus 0-II) and advanced-stage disease (Braak stages V-VI versus 0-II).

### Meta-analysis of genome-wide miRNA association analyses

To combine differential miRNA expression signatures across independent sample repositories, meta-analyses were conducted using the *metafor* R package^35^ (v5.0-1). Cohort-specific effect sizes and their corresponding standard errors were extracted for each miRNA. To guarantee analytical robustness and account for potential cross-cohort heterogeneity, both fixed-effect and random-effects models were systematically fitted using the maximum-likelihood estimation function. Combined effect sizes, pooled standard errors, and inter-cohort heterogeneity metrics were calculated across all contributing cohorts for both modeling frameworks, and their respective results were reported. Multiple testing corrections were applied globally using the Bonferroni method.

### miRNA target identification and tissue-specific filtering

To determine the downstream molecular pathways influenced by disease-associated post-transcriptional deregulation, validated and predicted gene targets of the miRNAs identified as statistically significant in the meta-analysis were retrieved using the *multiMiR* R package^36^ (v1.32.0; Database Version: 2.4.0 Updated: 2024-08-28). The human reference database was queried using the core retrieval function, extracting experimentally supported and computationally predicted interactions across eleven distinct specialized repositories, including miRecords^37^, miRTarBase^38^, TarBase^39^, DIANA-microT^40^, ElMMo^41^, MicroCosm^42^, miRanda^43^, miRDB^44^, PicTar^45^, PITA^46^, and TargetScan^47^. The underlying database configuration was locked to a fixed version utilizing the internal database version switch function to guarantee complete analytical reproducibility. To ensure biological relevance within the context of the human central nervous system, the resulting mRNA target pool was systematically filtered to retain only those transcripts with verified expression in cortical brain tissue across the UKBBN, BDR-1, PITT-ADRC, OPTIMA and ROSMAP total RNA sequencing datasets.

### Integration of miRNA-mRNA and miRNA-protein pairs

To map functionally relevant post-transcriptional regulatory units, miRNA-mRNA and miRNA-protein pairs were prioritized for downstream characterization when both molecular components within a pair displayed significant differential expression across the respective NFT Braak stage contrasts. Crucially, pairs were retained only when the target mRNA was Bonferroni significantly associated with Braak stage in the meta-analysis across the UKBBN, BDR-1, PITT-ADRC, OPTIMA, and ROSMAP datasets (*p*_adj <_ 0.05), or when the target protein showed a nominally significant association (*p* < 0.05) in the ROSMAP cohort. In addition, targets were required to exhibit an inverse direction of effect relative to their upstream miRNA regulator, consistent with the expected post-transcriptional silencing effect of miRNAs. Next, for each miRNA-mRNA and miRNA-protein pair in each individual dataset, linear regression models were fitted by designating target mRNA or protein expression levels as the continuous outcome and upstream miRNA abundance as the primary predictor and performing in parallel in each dataset. To isolate direct regulatory associations, these models were adjusted for clinical and technical factors including age at death, sex, tissue RIN, and cohort-specific technical covariates.

To generate consensus association metrics for the miRNA-mRNA pairs across the five cohorts, cohort-specific association estimates and their corresponding standard errors were pooled using an inverse-variance weighted random-effects meta-analysis framework and statistical significance was defined as an FDR-adjusted *p*-value (*p_adj)_* < 0.05, with the same significance threshold used for the miRNA-protein pairs.

### Causal mediation modeling of microRNA-associated tau pathology transitions

To assess whether miRNA expression mediated associations between transcriptomic or proteomic changes and tau pathology, mediation analyses were performed using the *mediation* R package^48^ (v4.5.1).

Analyses were restricted to miRNA-mRNA and miRNA-protein pairs identified in the preceding integrative analyses and models were adjusted for age at death and sex. Analyses used complete observations for each pair and required at least 20 samples and non-zero variance in mediator and outcome expression. Average causal mediation effects (ACME), average direct effects (ADE), total effects, and proportions mediated were estimated, with 95% confidence intervals obtained using nonparametric bootstrapping with 1,000 resamples. For mRNA outcomes, analyses were performed separately in the four PFC cohorts (ROSMAP, BDR, UKBBN, and PITT-ADRC); mediation was considered replicated when the ACME had an unadjusted *p* < 0.05 in at least three of the four cohorts with the same direction of effect. For protein outcomes, analyzed in ROSMAP, significance was defined as a Benjamini–Hochberg (BH) false discovery rate (FDR) *p*-value (*p*_adj)_ < 0.05 for the ACME. Significant mediation patterns were classified as predominantly miRNA-mediated when the ADE was not significant (unadjusted *p* ≥ 0.05), reinforcing when significant ACME and ADE estimates had the same direction, or opposing when they had opposite directions.

### Consensus gene co-expression network analysis and miRNA-module associations

To identify coordinated transcriptional programs associated with AD-related miRNAs, consensus weighted gene co-expression network analysis (WGCNA) was performed across the four PFC transcriptomic datasets (BDR-1, PITT-ADRC, UKBBN, and ROSMAP). mRNA expression matrices were residualized using linear models implemented in *limma*, adjusting for age at death, sex, RIN, sequencing batch, brain bank (where applicable), and the five estimated surrogate variables (SV1-SV5). Within each cohort, genes in the top 50% of the median absolute deviation (MAD) were retained, and only genes present across all four datasets after filtering (N = 3,180) were included for network construction.

The scale-free topology criterion was evaluated independently in each cohort using the pickSoftThreshold function in the *WGCNA* R package^29^ (v1.74), and a consensus soft-thresholding power of β = 6 was selected (**Supplementary Fig. 2**). Consensus co-expression modules were identified using the blockwiseConsensusModules function with a signed network, Pearson correlation, minimum module size of 30 genes, deepSplit = 2, and mergeCutHeight = 0.25. Module eigengenes were calculated separately for each cohort, and the unassigned grey module was excluded from downstream analyses.

Module eigengenes were correlated with expression levels of the Bonferroni-significant AD-associated miRNAs and Braak stage using Pearson correlation. Correlations were calculated separately for each cohort and module-miRNA associations were considered reproducible when they reached nominal significance (*p* < 0.05) with concordant directions of effect in at least three of the four cohorts. Mean correlation coefficients were subsequently calculated across cohorts for reproducibly associated module-miRNA pairs.

Genes from selected consensus modules were subjected to Gene Ontology over-representation analysis using clusterProfiler^49^ (v4.18.1). Redundant Gene Ontology terms were removed using the simplify function with Wang semantic similarity (cutoff = 0.7).

### Cell type-specific mapping of the target genes

Published single-nucleus RNA sequencing (snRNA-seq) differential expression data in association with NFT Braak stage from human prefrontal cortex in the ROSMAP cohort (Mathys *et al*.)^11^ were used to assign miRNA-mediated target genes identified in the bulk analysis to individual brain cell types. Cluster-level differential expression results were obtained for each cell type, and genes significantly associated with NFT pathology at a within-cell-type Bonferroni-adjusted *p*-value < 0.1 were retained. This threshold was selected to preserve sensitivity in the relatively small cell-type-specific gene sets and was used solely to define input gene sets for this secondary enrichment analysis.

All genes tested within each cell type served as the background for enrichment analyses. Enrichment of miRNA target genes among NFT-associated genes was assessed using one-sided hypergeometric tests, with odds ratios calculated from 2 × 2 contingency tables. Enrichment significance was defined as BH-adjusted FDR < 0.1.

### Cell-type-specific gene regulatory network analysis

To characterize cell-type-specific regulatory programs associated with AD, we constructed directed three-layer gene regulatory networks integrating transcription factors (TFs), miRNAs, and downstream target genes. Networks were assembled by integrating (i) experimentally validated TF-miRNA interactions from the TransmiR database^50^, (ii) replicated miRNA-target relationships identified through our cortical mediation analyses, and (iii) human brain cell-type-specific regulatory interactions from Brain Single-Cell Omics for PsychEncode (BrainSCOPE) gene regulatory networks^51^.

Networks were projected onto individual cortical cell types using snRNA-seq differential expression data obtained from Mathys *et al*. study^11^. Brain cell clusters were matched to their corresponding BrainSCOPE reference networks, and TFs were retained only if they were significantly differentially expressed within the corresponding cell type (*p_adj_* _<_ 0.1).

Regulatory interactions were filtered using the direction of disease-associated expression changes. TF-miRNA edges were retained as activating or repressive based on concordant or discordant changes in TF and miRNA expression, respectively. miRNA-target edges were retained only when target-gene expression changed in the opposite direction to the corresponding miRNA, consistent with canonical miRNA-mediated repression.

Directed networks were constructed using the *igraph* R package^52^ (v2.3.3) following removal of duplicate edges and self-loops. Network topology was characterized within each cell type using in-degree, out-degree, betweenness centrality, and PageRank. Similarity between cell-type networks was quantified using the pairwise Edge Jaccard Similarity Index, calculated as the number of shared directed edges divided by the total number of unique directed edges between each network pair. Potential intercellular signaling interactions were identified by intersecting network target genes with the human CellChatDB ligand-receptor database^53^.

### Genetic co-localization analysis of miRNA expression and AD risk loci

To investigate whether genetic regulation of AD-associated miRNAs overlapped with AD susceptibility loci, we performed cis-miQTL mapping followed by Bayesian co-localization analysis. Cis-miQTL analysis was performed using Matrix eQTL^54^ (v2.3) using linear regression. Models were adjusted for age, sex, RIN category, Braak stage, and the first three genotype-derived principal components to account for population structure.

Clinical- and batch-adjusted expression levels of the sixteen miRNAs associated with advanced AD pathology were combined across all seven postmortem miRNA datasets together with the corresponding imputed genotype data. Cis-miQTL analyses were performed within a ±1 Mb window centered on each miRNA precursor locus. Variants associated with miRNA expression at *p* < 1 × 10^-3^ were retained for downstream co-localization analyses.

Bayesian co-localization was performed using the *coloc* R package^55^ (v5.2.3) with the coloc.abf function to evaluate whether cis-miQTLs and AD susceptibility identified by the consensus meta-analysis of genome-wide association studies for AD^12^ shared a common causal variant. Evidence for co-localization was defined as a posterior probability for hypothesis 4 (PP4) >0.90, indicating a high probability that both traits are driven by the same underlying causal variant.

## Data availability

miRNA-seq and RNA-seq data and corresponding phenotypic data used in this study is available on the Zenodo Platform (DOI:10.5281/zenodo.22750742). Due to the sensitive nature of raw genotyping data this is available to the community under reasonable request and data sharing agreements. For the ROSMAP cohort, genotyping, miRNA-sequencing, mRNA-sequencing and proteomic data is available through the synapse platform (https://www.synapse.org/) under accessions syn3219045.

## Code availability

Analysis scripts used in this manuscript are available at https://github.com/Dementia-Systems-Biology/Cortical_miRNA_AD_Metaanalysis.

## Acknowledgements

We gratefully acknowledge the brain banks used in the sourcing of tissue, clinical, and neuropathological data used in this study. This includes the following brain banks: The Oxford Brain Bank (BDR-1, BDR-2, OPTIMA), LNDBB (BDR-1, BDR-2, LNDBB), The Newcastle Brain Tissue Resource (BDR-1, BDR-2, UKBBN), The South West Dementia Brain Bank (BDR-1, BDR-2), The Manchester Brain Bank (BDR-2, UKBBN), The Queen Square Brain Bank (UKBBN), The Edinburgh Brain Bank (UKBBN), The PITT-ADRC Brain Donation Program (PITT-ADRC) and The Rush Alzheimer’s Disease Center (ROSMAP). We thank the donors and their families for making this research possible. We thank Drs Aaron Jefferies, Karen Moore and Paul O’Neill at the University of Exeter sequencing service for help with the small RNA sequencing experiments for the UKBBN, BDR-1, BDR-2, PITT-ADRC and LNDBB datasets. We thank Drs Sören Franzenburg and Janina Fuß at the Institute for Molecular Biology at Kiel University for help with the small RNA sequencing experiments in the OPTIMA samples.

## Contributions

K.L. and E.P. conceived and designed the study and co-wrote the original manuscript. J.I., L.W., L.F.M., G.W., V.D. and A.R.S. performed laboratory experiments. E.P. performed the analyses. M.K., M.M.A., M.S., J.H., D.S., G.T., G.P., J.K., and V.L. contributed to data quality control and within-cohort analyses. A.T., M.S., D.v.d.H., T.M., B.C. and L.B. contributed to interpretation of findings. C.T., S.A-S, L.P. and J.M. provided brain tissue for analysis. L.B. contributed to study design and provided data. E.P., L.B., B.C. and K.L. were involved in funding acquisition. All authors reviewed and approved the final version of the manuscript.

## Ethics declarations

All human post-mortem tissue and associated data were obtained in accordance with the ethical and regulatory requirements of the respective brain bank, with informed consent provided. Ethical approval for our study was granted from the University of Exeter Medical School Research Ethics Committee (13/02/009). This study complied with all relevant regulations regarding the use of human study participants and was conducted in accordance with the criteria set by the Declaration of Helsinki.

## REFERENCES

1. Delay, C., Mandemakers, W. & Hebert, S.S. MicroRNAs in Alzheimer’s disease. Neurobiol Dis 46, 285–90 (2012).

2. Long, J.M., Ray, B. & Lahiri, D.K. MicroRNA-339-5p down-regulates protein expression of beta-site amyloid precursor protein-cleaving enzyme 1 (BACE1) in human primary brain cultures and is reduced in brain tissue specimens of Alzheimer disease subjects. J Biol Chem 289, 5184–98 (2014).

3. Smith, P.Y. et al. miR-132/212 deficiency impairs tau metabolism and promotes pathological aggregation in vivo. Hum Mol Genet 24, 6721–35 (2015).

4. Liu, W. et al. The neuroinflammatory role of microRNAs in Alzheimer’s disease: pathological insights to therapeutic potential. Mol Cell Biochem 480, 2689–2706 (2025).

5. Kumar, S. & Reddy, P.H. The role of synaptic microRNAs in Alzheimer’s disease. Biochim Biophys Acta Mol Basis Dis 1866, 165937 (2020).

6. Salta, E. & De Strooper, B. Noncoding RNAs in neurodegeneration. Nat Rev Neurosci 18, 627–640 (2017).

7. Salta, E., Sierksma, A., Vanden Eynden, E. & De Strooper, B. miR-132 loss de-represses ITPKB and aggravates amyloid and TAU pathology in Alzheimer’s brain. EMBO Mol Med 8, 1005–18 (2016).

8. Dobricic, V. et al. Differential microRNA expression analyses across two brain regions in Alzheimer’s disease. Transl Psychiatry 12, 352 (2022).

9. Li, Q.S. & Cai, D. Integrated miRNA-Seq and mRNA-Seq Study to Identify miRNAs Associated With Alzheimer’s Disease Using Post-mortem Brain Tissue Samples. Front Neurosci 15, 620899 (2021).

10. Walgrave, H., Zhou, L., De Strooper, B. & Salta, E. The promise of microRNA-based therapies in Alzheimer’s disease: challenges and perspectives. Mol Neurodegener 16, 76 (2021).

11. Mathys, H. et al. Single-cell multiregion dissection of Alzheimer’s disease. Nature 632, 858–868 (2024).

12. Bellenguez, C. et al. Consensus meta-analysis of genome-wide association studies for Alzheimer’s disease and related dementias. Nature Genetics 58, 1214–1225 (2026).

13. Lau, P. et al. Alteration of the microRNA network during the progression of Alzheimer’s disease. EMBO Mol Med 5, 1613–34 (2013).

14. Wong, H.K. et al. De-repression of FOXO3a death axis by microRNA-132 and -212 causes neuronal apoptosis in Alzheimer’s disease. Hum Mol Genet 22, 3077–92 (2013).

15. Magill, S.T. et al. microRNA-132 regulates dendritic growth and arborization of newborn neurons in the adult hippocampus. Proc Natl Acad Sci U S A 107, 20382–7 (2010).

16. Hernandez-Rapp, J. et al. microRNA-132/212 deficiency enhances Abeta production and senile plaque deposition in Alzheimer’s disease triple transgenic mice. Sci Rep 6, 30953 (2016).

17. El Fatimy, R., et al. MicroRNA-132 provides neuroprotection for tauopathies via multiple signaling pathways. Acta Neuropathol 136, 537–555 (2018).

18. Walgrave, H. et al. Restoring miR-132 expression rescues adult hippocampal neurogenesis and memory deficits in Alzheimer’s disease. Cell Stem Cell 28, 1805–1821 e8 (2021).

19. Johnson, E.C.B. et al. Large-scale deep multi-layer analysis of Alzheimer’s disease brain reveals strong proteomic disease-related changes not observed at the RNA level. Nat Neurosci 25, 213–225 (2022).

20. Morabito, S. et al. Single-nucleus chromatin accessibility and transcriptomic characterization of Alzheimer’s disease. Nat Genet 53, 1143–1155 (2021).

21. Sadick, J.S., O’Dea, M.R., Hasel, P., Dykstra, T., Faustin, A. & Liddelow, S.A. Astrocytes and oligodendrocytes undergo subtype-specific transcriptional changes in Alzheimer’s disease. Neuron 110, 1788–1805 e10 (2022).

22. Grubman, A. et al. A single-cell atlas of entorhinal cortex from individuals with Alzheimer’s disease reveals cell-type-specific gene expression regulation. Nat Neurosci 22, 2087–2097 (2019).

23. Nagaraj, S. et al. Differentially expressed miRNAs in the temporal cortex of Alzheimer’s disease patients and their association to tau pathology. Commun Biol 9, 253 (2026).

24. Kouhsar, M. et al. A brain DNA co-methylation network analysis of psychosis in Alzheimer’s disease. Alzheimers Dement 21, e14501 (2025).

25. Pegoraro, G. et al. Systemic infections alter cortical transcriptional signatures in Alzheimer’s disease. Neurobiology of Aging 167, 155–167 (2026).

26. Dobricic, V. et al. Common signatures of differential microRNA expression in Parkinson’s and Alzheimer’s disease brains. Brain Commun 4, fcac274 (2022).

27. Vattathil, S.M. et al. Mapping the microRNA landscape in the older adult brain and its genetic contribution to neuropsychiatric conditions. Nat Aging 5, 306–319 (2025).

28. Friedlander, M.R., Mackowiak, S.D., Li, N., Chen, W. & Rajewsky, N. miRDeep2 accurately identifies known and hundreds of novel microRNA genes in seven animal clades. Nucleic Acids Res 40, 37–52 (2012).

29. Langfelder, P. & Horvath, S. WGCNA: an R package for weighted correlation network analysis. Bmc Bioinformatics 9(2008).

30. Chen, Y., Chen, L., Lun, A.T.L., Baldoni, P.L. & Smyth, G.K. edgeR v4: powerful differential analysis of sequencing data with expanded functionality and improved support for small counts and larger datasets. Nucleic Acids Res 53(2025).

31. Leek, J.T., Johnson, W.E., Parker, H.S., Jaffe, A.E. & Storey, J.D. The sva package for removing batch effects and other unwanted variation in high-throughput experiments. Bioinformatics 28, 882–3 (2012).

32. Manichaikul, A., Mychaleckyj, J.C., Rich, S.S., Daly, K., Sale, M. & Chen, W.M. Robust relationship inference in genome-wide association studies. Bioinformatics 26, 2867–73 (2010).

33. Purcell, S. et al. PLINK: a tool set for whole-genome association and population-based linkage analyses. Am J Hum Genet 81, 559–75 (2007).

34. Ritchie, M.E. et al. limma powers differential expression analyses for RNA-sequencing and microarray studies. Nucleic Acids Res 43, e47 (2015).

35. Viechtbauer, W. Conducting Meta-Analyses in R with the metafor Package. Journal of Statistical Software 36, 1–48 (2010).

36. Ru, Y. et al. The multiMiR R package and database: integration of microRNA-target interactions along with their disease and drug associations. Nucleic Acids Res 42, e133 (2014).

37. Xiao, F., Zuo, Z., Cai, G., Kang, S., Gao, X. & Li, T. miRecords: an integrated resource for microRNA-target interactions. Nucleic Acids Res 37, D105–10 (2009).

38. Hsu, S.D. et al. miRTarBase: a database curates experimentally validated microRNA-target interactions. Nucleic Acids Res 39, D163–9 (2011).

39. Vergoulis, T. et al. TarBase 6.0: capturing the exponential growth of miRNA targets with experimental support. Nucleic Acids Res 40, D222–9 (2012).

40. Maragkakis, M. et al. DIANA-microT Web server upgrade supports Fly and Worm miRNA target prediction and bibliographic miRNA to disease association. Nucleic Acids Res 39, W145–8 (2011).

41. Gaidatzis, D., van Nimwegen, E., Hausser, J. & Zavolan, M. Inference of miRNA targets using evolutionary conservation and pathway analysis. BMC Bioinformatics 8, 69 (2007).

42. Griffiths-Jones, S., Saini, H.K., van Dongen, S. & Enright, A.J. miRBase: tools for microRNA genomics. Nucleic Acids Res 36, D154–8 (2008).

43. Betel, D., Wilson, M., Gabow, A., Marks, D.S. & Sander, C. The microRNA.org resource: targets and expression. Nucleic Acids Res 36, D149–53 (2008).

44. Wang, X. miRDB: a microRNA target prediction and functional annotation database with a wiki interface. RNA 14, 1012–7 (2008).

45. Krek, A. et al. Combinatorial microRNA target predictions. Nat Genet 37, 495–500 (2005).

46. Kertesz, M., Iovino, N., Unnerstall, U., Gaul, U. & Segal, E. The role of site accessibility in microRNA target recognition. Nat Genet 39, 1278–84 (2007).

47. Grimson, A., Farh, K.K., Johnston, W.K., Garrett-Engele, P., Lim, L.P. & Bartel, D.P. MicroRNA targeting specificity in mammals: determinants beyond seed pairing. Mol Cell 27, 91–105 (2007).

48. Tingley, D., Yamamoto, T., Hirose, K., Keele, L. & Imai, K. mediation: R Package for Causal Mediation Analysis. Journal of Statistical Software 59(2014).

49. Xu, S. et al. Using clusterProfiler to characterize multiomics data. Nat Protoc 19, 3292–3320 (2024).

50. Tong, Z., Cui, Q., Wang, J. & Zhou, Y. TransmiR v2.0: an updated transcription factor-microRNA regulation database. Nucleic Acids Res 47, D253–D258 (2019).

51. Emani, P.S. et al. Single-cell genomics and regulatory networks for 388 human brains. Science 384, eadi5199 (2024).

52. Antonov, M. et al. igraph 1.0 enables fast and robust network analysis across programming languages. Plos One 21(2026).

53. Jin, S., Plikus, M.V. & Nie, Q. CellChat for systematic analysis of cell-cell communication from single-cell transcriptomics. Nat Protoc 20, 180–219 (2025).

54. Shabalin, A.A. Matrix eQTL: ultra fast eQTL analysis via large matrix operations. Bioinformatics 28, 1353–8 (2012).

55. Wallace, C. A more accurate method for colocalisation analysis allowing for multiple causal variants. PLoS Genet 17, e1009440 (2021).

