## Supplementary Figures 1-2 for "A large-scale multi-omics study of cortical miRNA dysregulation in Alzheimer’s disease"

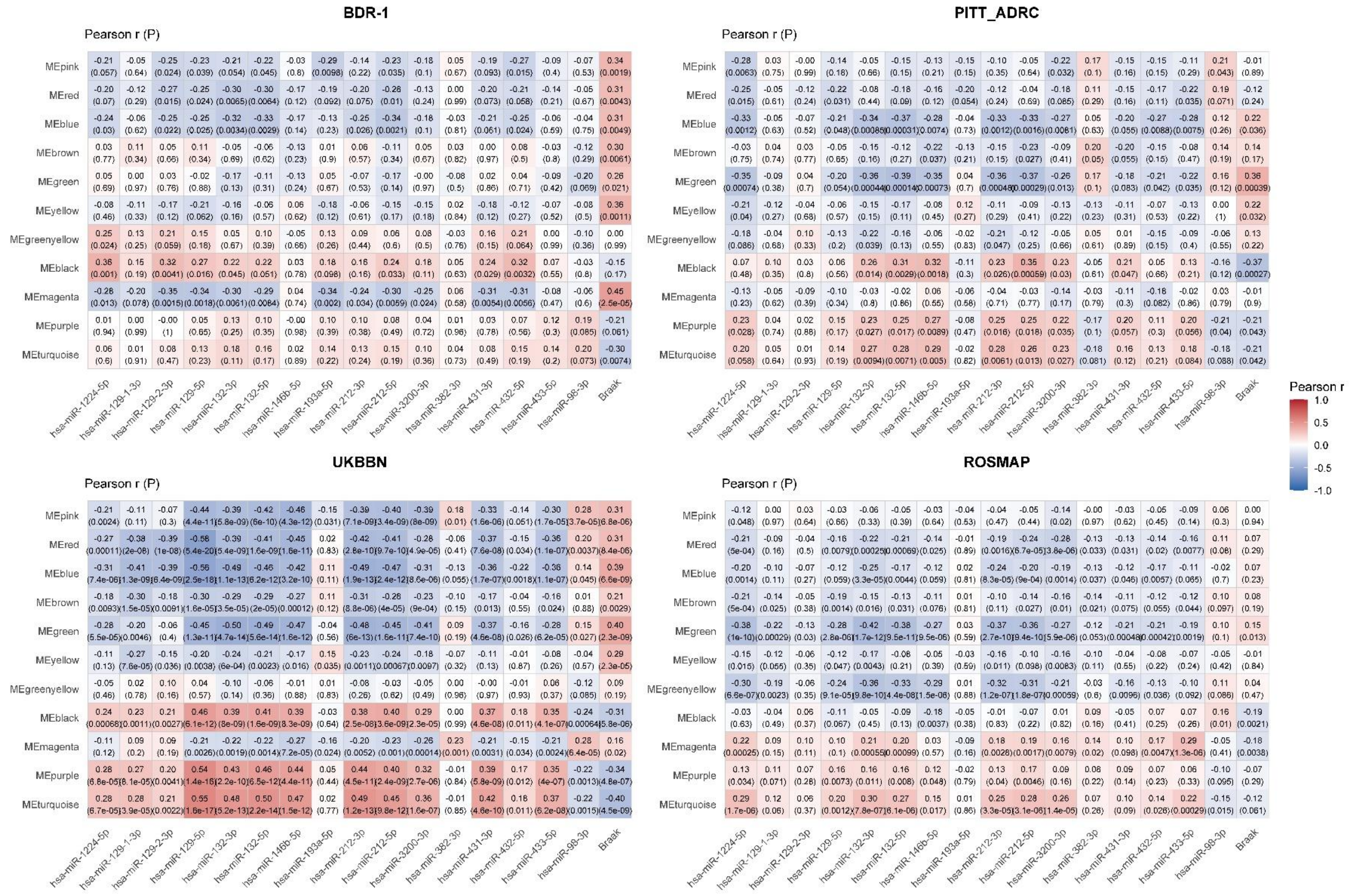


**Supplementary Fig. 1: Cohort-specific module–trait relationships.** Heatmaps showing Pearson correlations between consensus WGCNA module eigengenes and AD-associated miRNAs across the four independent prefrontal cortex (PFC) RNA-sequencing datasets (BDR-1, PITT-ADRC, UKBBN, and ROSMAP). Module eigengenes were correlated with the expression levels of Bonferroni-significant miRNAs identified in the cross-cortical meta-analysis, together with Braak neurofibrillary tangle (NFT) stage. Values within tiles indicate Pearson correlation coefficients (*r*); asterisks denote FDR-significant associations within each cohort. Red indicates positive correlations and blue indicates negative correlations.

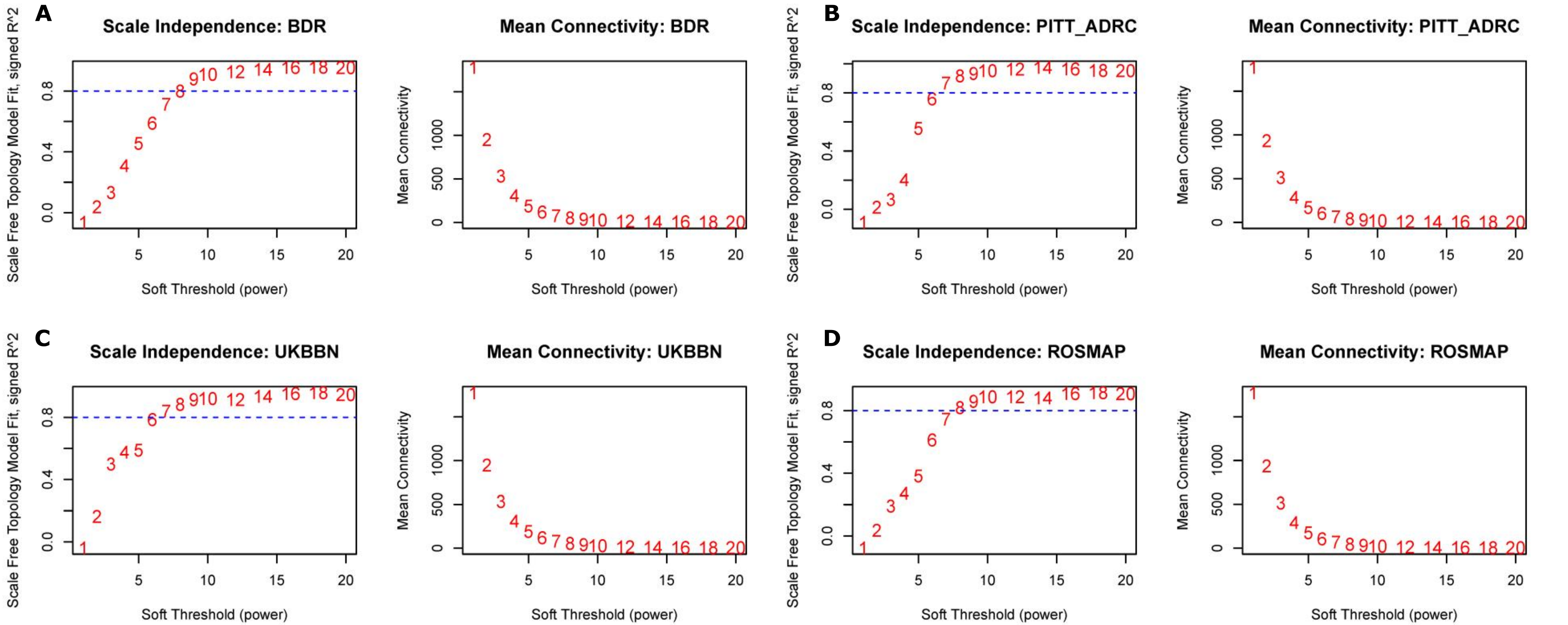


**Supplementary Fig. 2: Selection of the soft-thresholding power for consensus WGCNA.** Scale-free topology model fit (left) and mean connectivity (right) across a range of candidate soft-thresholding powers for each RNA-sequencing cohort (**a** BDR-1, **b** PITT-ADRC, **c** UKBBN, and **d** ROSMAP). Numbers indicate the tested soft-thresholding power. The dashed blue line denotes the conventional threshold for an acceptable signed scale-free topology fit (0.80). The final soft-thresholding power used for consensus network construction was selected as a compromise across cohorts to achieve an approximately scale-free topology while maintaining sufficient network connectivity.
